# Goiter parenchyma, blood vessels and lymphatics contain Staphylococcus epidermidis - saprophyte or pathogen?

**DOI:** 10.1101/719732

**Authors:** Sergiusz Durowicz, Marzanna Zaleska, Waldemar L. Olszewski, Ewa Stelmach, Katarzyna Piskorska, Ewa Swoboda-Kopeć, Wiesław Tarnowski

## Abstract

**Background:** Goiter in its various clinical and histopathological forms is accompanied by an inflammatory process requiring intensive therapy. The thyroid gland is an organ specifically exposed to the microbial environment due to its close location to the mouth microbiome. A number of bacterial phenotypes has been detected in the inflamed thyroid gland. A question raises as to whether bacteria have not already been present in the thyroid gland before the clinical symptoms of goiter became evident.

**Aim:** To answer the questions: a) do the goiter tissue structures contain bacteria, b) if so, which bacterial phenotypes can be identified, c) what are the genetic similarities of the thyroid and periodontal bacterial strains.

**Material and methods:** Studies were carried out in 60 patients with the non-toxic multinodular goiter in 40 cases, toxic multinodular goiter in 10, single adenoma in 3, Hashimoto’s disease with nodular changes in 4 and recurrent thyroid disease in 3. Tissue fragments harvested during surgery were placed on Columbia blood agar base enriched with 5% defibrinated sheep blood. In this method bacteria present in the tissue slowly proliferate in their in vivo transferred to ex-vivo environment, crawl out and form the on-plate colonies. It enables detection of single bacteria usually difficult in a standard planktonic culture.

**Results:** a) Coagulase-negative Staphylococci were shown growing on culture plates in above 50% of thyroid parenchyma, veins, arteries and adjacent lymphoid tissue specimens, b) tissue-originating colony-forming bacteria appeared on plates on day 3, but in some as late as after 12-21 days, c) all isolates were sensitive to the basic antibiotics, d) bacterial thyroid and oral DNA tests showed similarities indicating possibility of the oral origin, e) the on-plate time-prolonged cultures showed shrinking of the colonies and upon adding liquid medium formed the small variant colonies.

**Conclusions:** Thyroid gland tissues contained in above 50% of specimens the coagulase-negative Staphylococci. Over 88% similarity of the genetic pattern of Staphylococcus epidermidis strain from tooth, oropharyngeal and thyroid tissues, estimated with PCR MP technique, suggested their periodontium origin.

## Introduction

Goiter in its various clinical and histopathological forms is often an inflammatory process requiring intensive therapy. The thyroid gland is an organ specifically exposed to the microbial environment due to its close location to the mouth microbiome. A number of publications has been reported on a spectrum of bacterial strains identified in the thyroid in cases with thyroiditis [1–3]. Despite the name “chronic lymphocytic”, in Hashimoto’s disease the follicular structures are infiltrated not only by lymphocytes (72%) but also eosinophils (48%) and neutrophils (26%) [4]. Neutrophil infiltrates in the histopathological pictures may indicate the presence of bacteria in the thyroid parenchyma. Viral infections are also frequently cited as a major environmental factor involved in subacute thyroiditis and autoimmune thyroid diseases [5].

However, it remains to determine whether bacteria and viruses are responsible for thyroid diseases or they are just innocent bystanders. Moreover, it should be known whether they might be responsible for inflammation in the gland bed and skin flap after surgery.

The thyroid gland is presumed to be evolutionarily resistant to infection due to its abundant vascularity, lymphatic drainage, the presence of iodine and hydrogen peroxide in the tissue, and because of its encapsulation. However, a number of bacterial phenotypes has been detected in the inflamed thyroid gland as Staphylococcus aureus, pyogenes, and epidermidis, and Streptococcus pneumoniae. Occasionally other aerobic organisms as Klebsiella sp, Haemophilus influenza, Streptococcus viridans, Eikenella corrodens, Enterobacteriaceae, and Salmonella spp were identified [6–8].

Another problem is the relatively uncommon but difficult for control complication of thyroidectomy as wound healing with inflammation, skin flap swelling and further excessive scarring. The incidence of seroma after thyroidectomy has been reported by some authors to be between 0.3% and 7%. It increases with the extent of surgery, with a higher incidence reported after bilateral procedures or after the removal of large substernal goiters [9–10]. The postthyroidectomy skin sinus develops in 0.088% and albeit rare is difficult for treatment [11].

The mechanism of colonization of the thyroid by microbes is still unclear. Presumably, bacteria can reach it via lymphatics from areas of the oropharyngeal infection or as hematogenous spread from a remote infection. There is an increasing body of evidence for translocations of microbes from tissues as mouth, gut or skin to the peripheral tissues [12–14].

A question raises as to whether bacteria have not already been present in the thyroid gland before the clinical symptoms of thyroiditis became evident. A large body of literature is now available on the presence of microbes in the apparently “sterile” tissues as e.g. female reproductive organs, brain, bones and limb vascular bundles [15–22].

Tissue specimens may contain single microbes dwelling in the tissue structure, but they are not easily detected in the specimens using the standard liquid bacteriological culture media. We have found that in case the harvested tissue fragments during surgery are placed for some days on a culture plate, bacteria may slowly proliferate in their cultured tissue, crawl out and form colonies. Moreover, this technique allows observation of bacteria forming colonies around as well as on various tissue topographical fragments as gland cells, blood vessels, nerves etc. what is not available using cultures of a homogenized specimen In this study we harvested in a group of patients with diagnosed goiter thyroid fragments, arterial and vein walls, and draining lymph nodes, cultured them on-plates and observed daily the emerging bacterial colonies. The colonies could be observed already in a few days, in some cases as late as after 2-3 weeks. At prolonged culture periods formation of a small colony variant - dormancy state bacteria called “persisters” could also be seen. The genotyping of the tissue on-plate grown bacteria from thyroid and dental sacs was carried out to indicate their possible similarities.

The questions we posed were: a) do the goiter tissue structures contain bacteria, b) if so, which bacterial phenotypes can be identified, c) what are the genetic similarities of the thyroid and periodontal bacterial strains,

## Material and methods

### Patients

Studies were carried out in 60 patients admitted to the surgical department for thyroid surgery according to the order they showed up for therapy. There were 55 females and 5 males, median age was 54 years (range 21 to 78). The preoperative clinical diagnosis was in 40 cases the non-toxic multinodular goiter, in 10 the toxic multinodular goiter, in 3 single adenoma, in 4 Hashimoto’s disease with nodular changes and in 3 recurrent thyroid disease with discomfort causing tracheal compression. Excluded were subjects with acute or chronic infection at remote sites and treated with antibiotics over the last 3 months. All patients signed consent for pharmacological and surgical treatment. The therapy protocol included the routine hospital procedures including bacteriology. Patients were duly informed on details of the study and gave consent for publishing the data. The study was approved by the ethics committee of the Postgraduate Medical School, Warsaw. This observational study was not a subject requiring EU registration.

### Surgical treatment

Surgical procedures included 37 total extracapsular thyroidectomies, 12 Dunhill operations, 8 hemi-thyroidectomies and 3 reoperations for benign recurrent condition.

### Collection of samples for bacteriology

#### Thyroid tissue fragments

Specimens were collected in the operating room under routine strictly sterile conditions. The operating room air and equipment control bacteriology using sedimentation and imprint tests was done. No growth was detected. The operation field skin was disinfected with Kodan Tinktur forte (Schülke & Mayr, Germany). All instruments and material used for collection of specimens were tested for sterility prior to their use in the study. Fragments of: a) incision skin, b) thyroid parenchyma, c) thyroid tissue artery, (d) thyroid tissue vein, and (e) pre-tracheal thyroid draining lymph nodes were harvested. Each sample was taken using a separate sterile instrument. Tissue specimens were placed on Columbia agar with sheep blood plate and cultured for at 37°C for up to 30 days. Once bacterial colonies appeared on or around the specimen a swab smear was taken and transferred to the transport media (Transwab MWE, UK) for the phenotype identification.

#### The oral vestibule gingival swabs

15 patients were swabbed for bacteriological identification from the oral vestibule, close to gingival pockets.

### Identification of bacterial strains

Specimens were checked daily for appearance of colonies on and around the tissue specimens and their size and color were evaluated.

The following media were used: Columbia blood agar base enriched with 5% sterile defibrinated sheep blood, MacConkey’s agar, Chapman’s agar, Sabouraud’s agar (malt agar), and brain heart infusion (BHI) (all from Difco, Detroit, MI). Swabs were taken from the on-plate colonies for further identifications. The cultures were incubated at 37°C and examined at 24 and 48 hr for aerobic bacterial growth. After 48 hr of incubation, BHI cultures were transferred into blood agar, and examined every day for up to 10 days and, if positive, inoculated onto blood agar slants. After 10 days, all negative cultures were transferred to blood agar. In cases where there was no aerobic growth, additional cultures for anaerobic growth were established. Isolates were identified by standard procedures using the Analytical Profile Identification (API) System (Biomerieux). Aerobic Cocci of the Micrococcaccae family were identified using the API- Staph system. Bacteria of Streptococcaceae family were identified with API 20 Strep.

### Antibiotic sensitivity

The sensitivity of isolated bacterial strains to antibiotics was examined using the ATB system (Biomerieux, Paris, France). Analysis of the antibiotic sensitivity was performed using the ATB-Plus reader (Biomerieux, Paris, France)

### Polymerase Chain Reaction Melting Profiles (PCR MP)

In order to study whether the thyroid tissue bacteria, identified previously with culturing methods, may originate from the oral cavity, the DNA patterns of isolated strains were compared. Genomic bacterial DNA isolation was performed with EurX Genomic DNA Purification Kit, according to the producer’s guidelines (EurX, Gdańsk, Poland). The DNA elution was performed with Tris-EDTA buffer and stored for further analysis at - 20°C.The genetic relatedness analysis was performed with PCR MP kit (DNA Gdańsk II, Poland), according to the manufacturer’s guidelines. In first step, the genomic DNA was digested by the restriction enzyme HindIII. In next step, the restriction fragments were ligated with an oligonucleotide adapter. Finally, during PCR the DNA fragments were amplified. Reactions of amplification were conducted in the DNA Engine thermocycler (BioRad, Hercules, Calif, United States), in the following conditions: the initial denaturation 72°C - 1 min, next 23 cycles: 72°C - 1 min, 81°C - 30 s, 72°C 2 min 30 s, and final elongation 72°C - 5 min. The products of the analysis were separated eletrophoretically in 2 % agarose gel with the addition of ethidium bromide and visualized by illumination with the ultraviolet light. The weight size of the obtained PCR products was compared to the molecular-weight size marker. The analysis of the results was performed using the GeneTools program (Syngene, Cambridge, United Kingdom).

### Statistical evaluation

Descriptive statistics are presented as the number, mean, standard deviation, minimum and maximum for continuous variables and as the number and percentage for categorical variables. For evaluation of statistical differences between the numerical prevalence of bacterial isolates the Chi-square test was applied. The analysis was performed using SPSS 25.0 statistical software.

## Results

### Tissue specimen on-plate bacterial culturing

Bacterial growth was observed on skin fragments in 26.9%, thyroid tissue in 61.7 %, thyroid arteries in 43.1%, veins in 45.7%, and lymphoid tissue in 52.7% (Table 1).

**Table 1.** Frequency of isolates from the specimens of thyroid tissues cultures on blood plates

|  | Skin | Vein | Artery | Thyroid | Lymph node |
| --- | --- | --- | --- | --- | --- |
| Number of taken specimens | 52 | 59 | 51 | 60 | 55 |
| Bacteria Strain |  |  |  |  |  |
| Staph epidermidis | 10 | 15 | 12 | 22 | 20 |
| Staph capitis | 3 | 5 | 2 | 7 | 2 |
| Staph saprophyticus |  |  | 2 | 1 | 1 |
| Staph aureus |  | 1 | 1 |  |  |
| Staph caprae |  |  |  | 1 |  |
| Staph xylosus |  |  |  |  | 1 |
| Staph warneri |  | 2 | 2 | 1 | 2 |
| Staph lugdunensis |  | 2 |  | 2 | 1 |
| Staph chromogenes |  | 1 |  | 1 |  |
| Staph haemolyticus |  | 1 |  |  | 1 |
| Total Staphylococci | 13/52 | 27/59 | 19/51 | 35/60 | 28/55 |
| % of growth | 25,0<sub>a</sub> | 45,7<sub>ab</sub> | 37,2<sub>ab</sub> | 58,3<sub>b</sub> | 50,9<sub>ab</sub> |
| $X^2(4) = 14,66; p = 0,005; V = 0,23$ | | | | | |
| Micrococcus luteus |  |  | 2 | 1 | 1 |
| Aerococcus viridans |  |  | 1 | 1 |  |
| Acinetobacter calcoaceticus var lwoffii | 1 |  |  |  |  |
| Total Bacteria | 14/52 | 27/59 | 22/51 | 37/60 | 29/55 |
| % of growth | 26,9<sub>a)</sub> | 45,7<sub>ab)</sub> | 43,1<sub>ab)</sub> | 61,7<sub>b)</sub> | 52,7<sub>ab)</sub> |
| $X^2(4) = 14,70; p = 0,005; V = 0,23$ | | | | | |

Statistical significance between a) and b).

The first colonies were identified optically on the specimen surface and around it from 3 to over 12 days (mean 4.3 days) (Fig. 1). Occasionally, first colonies appeared after 3 weeks. On continuation of culture the confluent colonies were formed adjacent to the specimens proving their origin from the tissue (Fig.2–4). In some cases small colonies could also were identified on specimens’ surface (Fig. 5). This may indicate that the natural tissue provides nutrients for bacterial proliferation and the plate culture medium is only secondary to it. There were many small colonies spreading around the culture plate upon supplementing with liquid medium (Fig. 6). These colonies belonged most likely to the so called persister strains. Interestingly, Bacterial colonies observed on plate for 2-3 weeks changed their spread from confluent to single colonies and didn’t proliferate (Fig. 3a and 4a).

**Fig 1.**
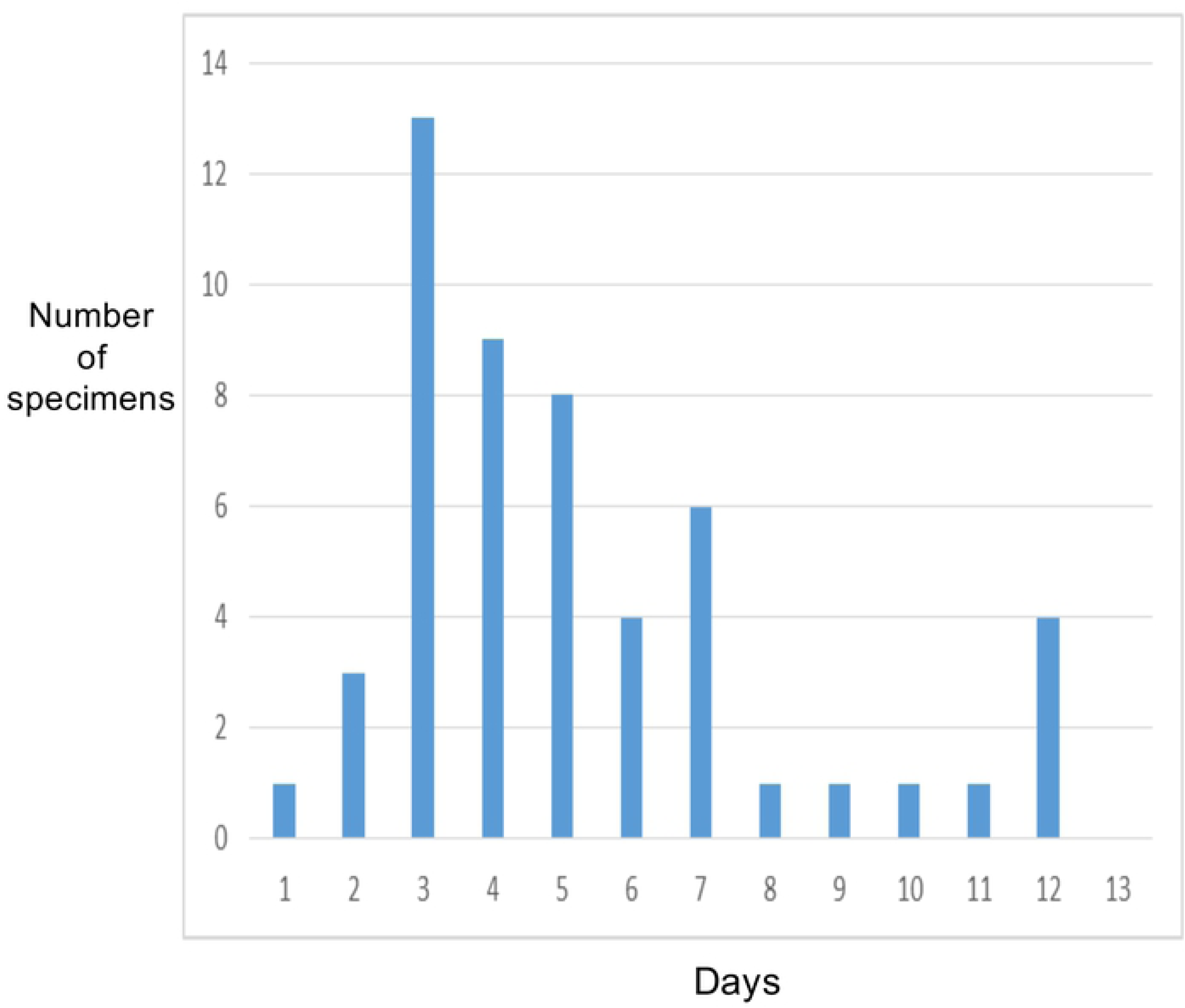
Schematic presentation of days of appearance of first bacterial colonies around and on the on-plate cultured specimens of the thyroid tissue.

**Fig 2.**
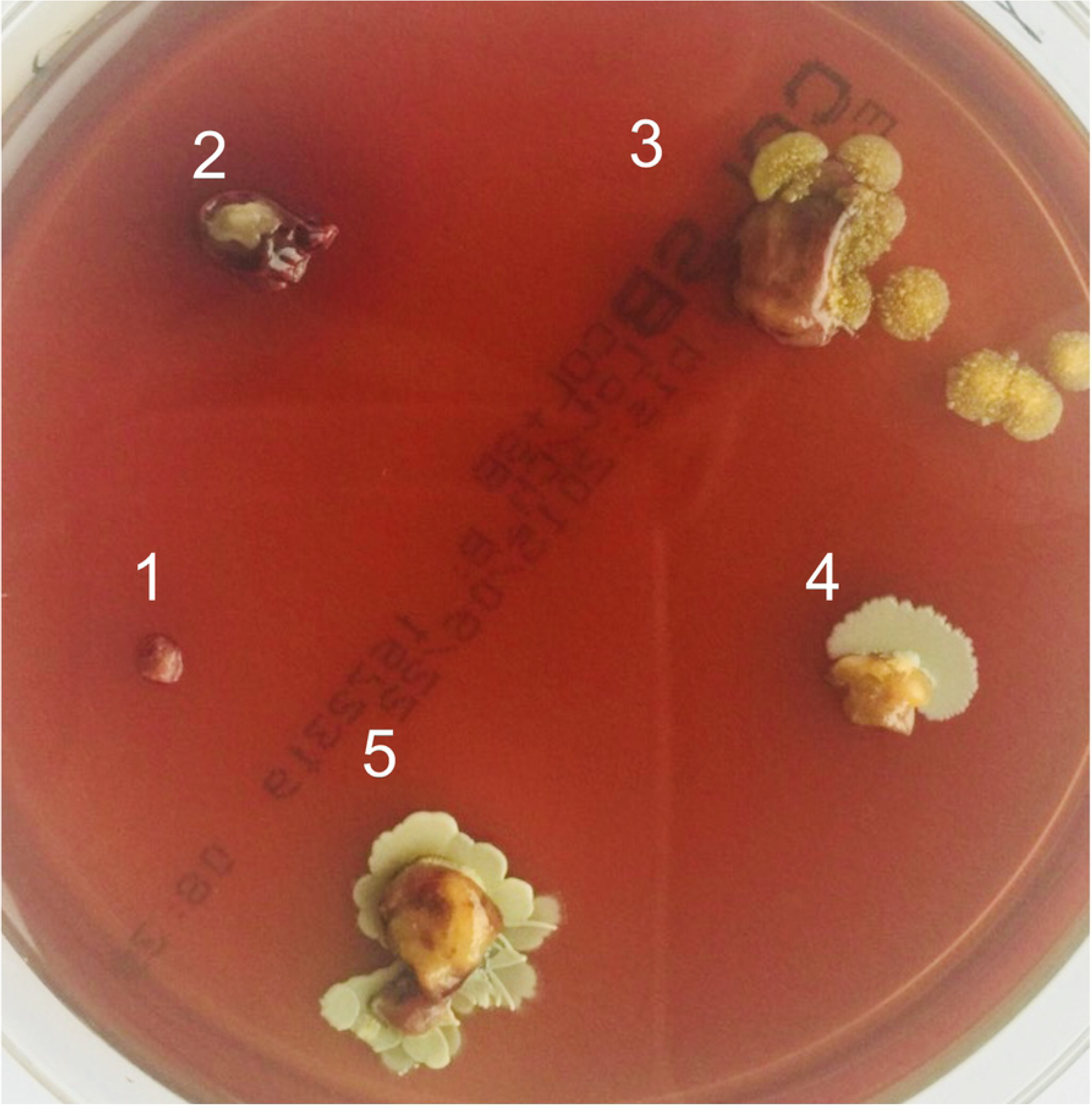
Thyroid tissue specimens on a culture plate one week after harvesting. 1. skin. 2. thyroid tissue, 3. vein, 4. artery, 5. lymphoid tissue. Bacterial colonies around thyroid structures but not skin.

**Fig 3.**
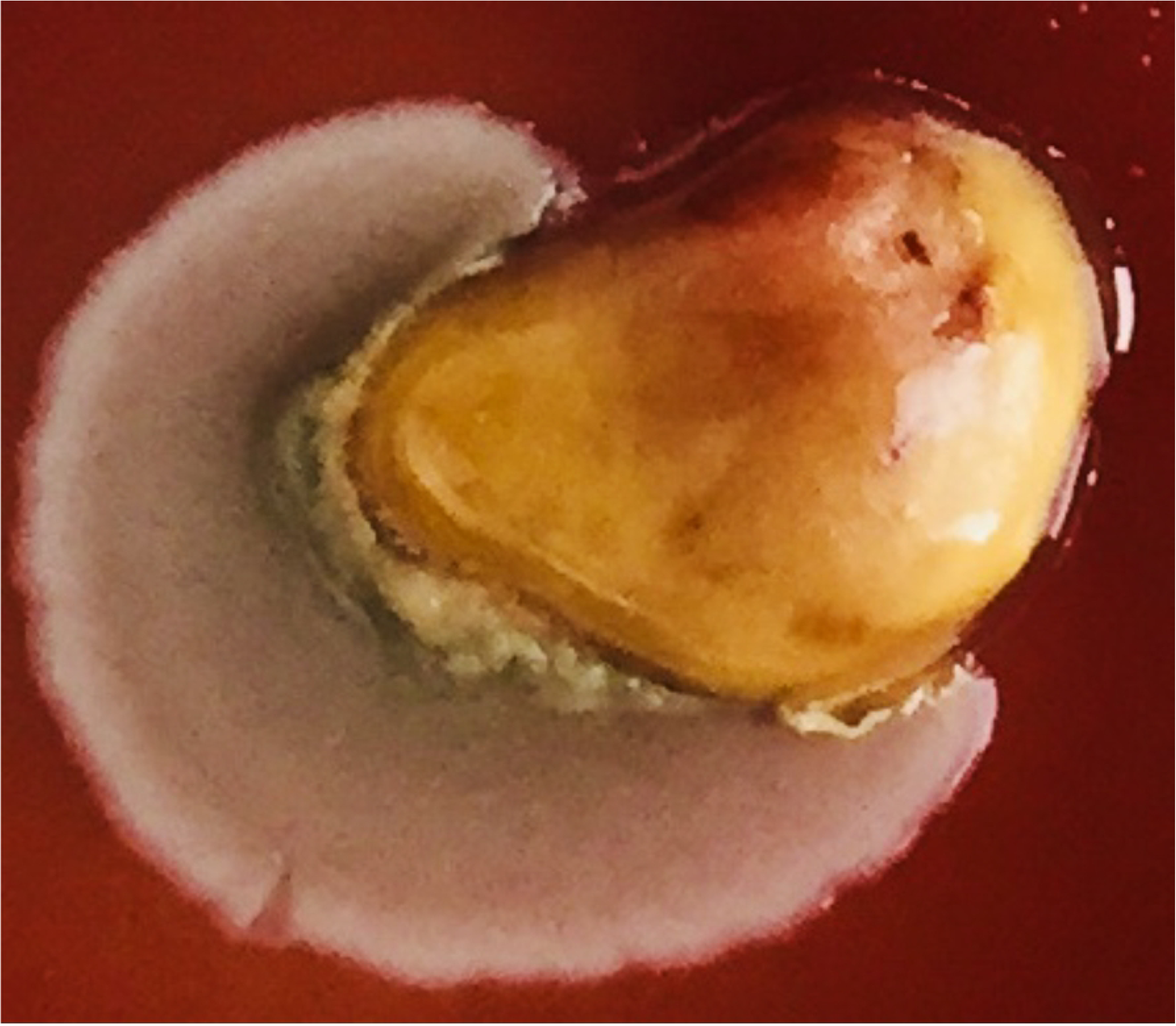
Thyroid specimen 7 days of culturing. A confluent spot of colonies by microbes migrating from the bottom part of tissue.

**Fig 3a.**
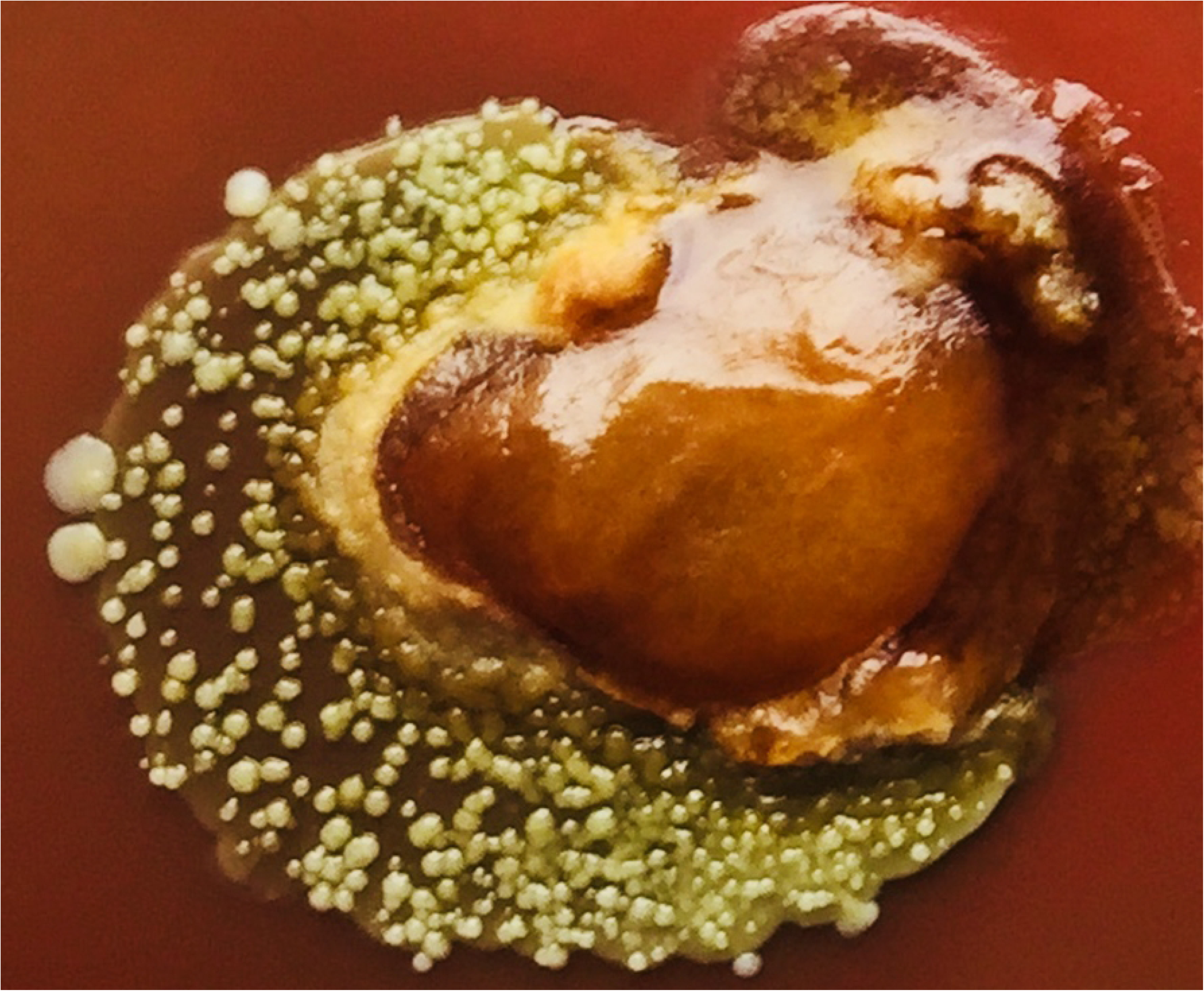
The same specimen as on Fig. 3 three weeks on culture. showing Multiple single colonies of various size differing from the early culture confluent growth. Three of them at the periphery are enlarged and more whitish.

**Fig 4.**
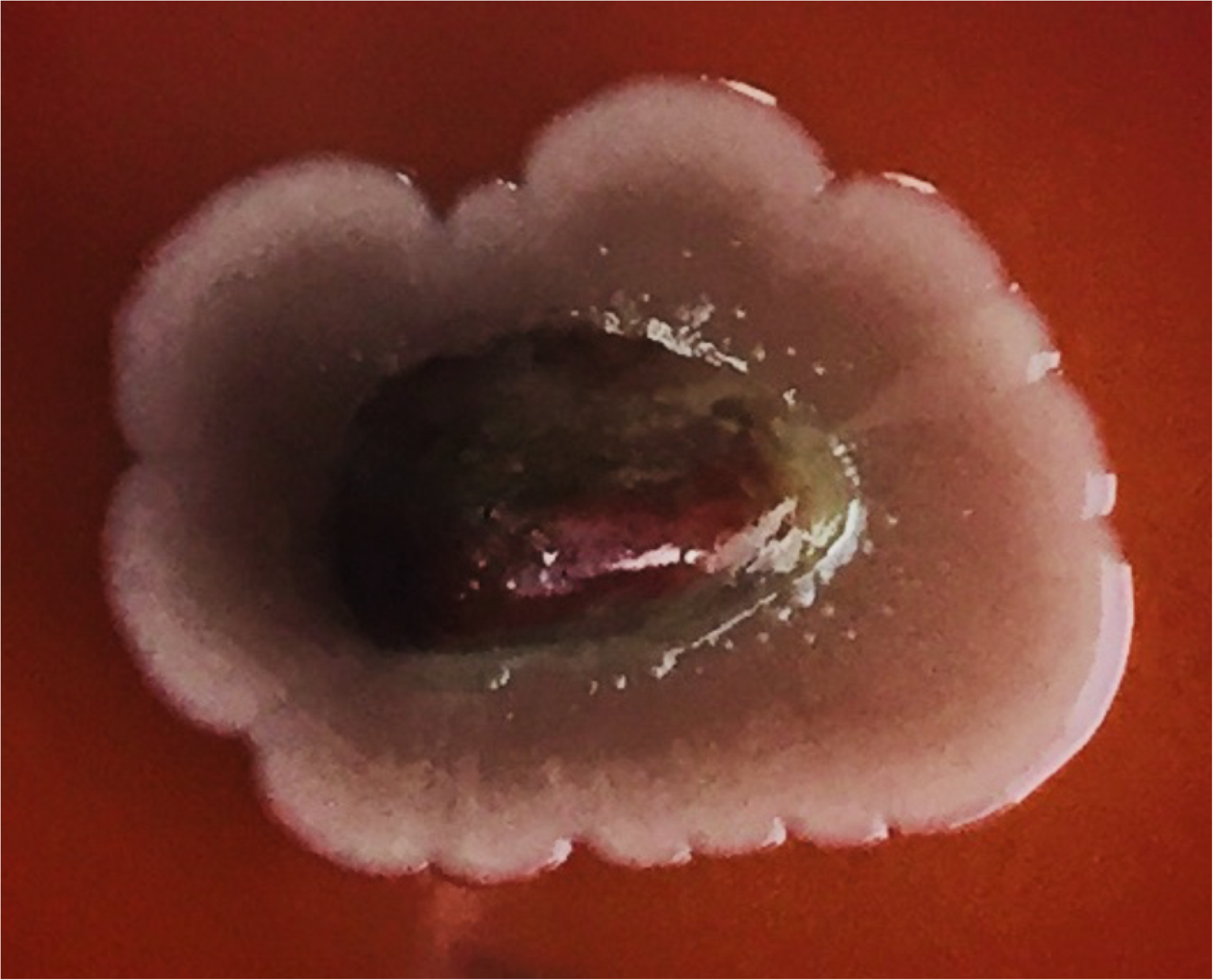
Another thyroid specimen on a culture plate five days after harvesting. Specimen surrounded by confluent colonies migrating from the tissue.

**Fig 4a.**
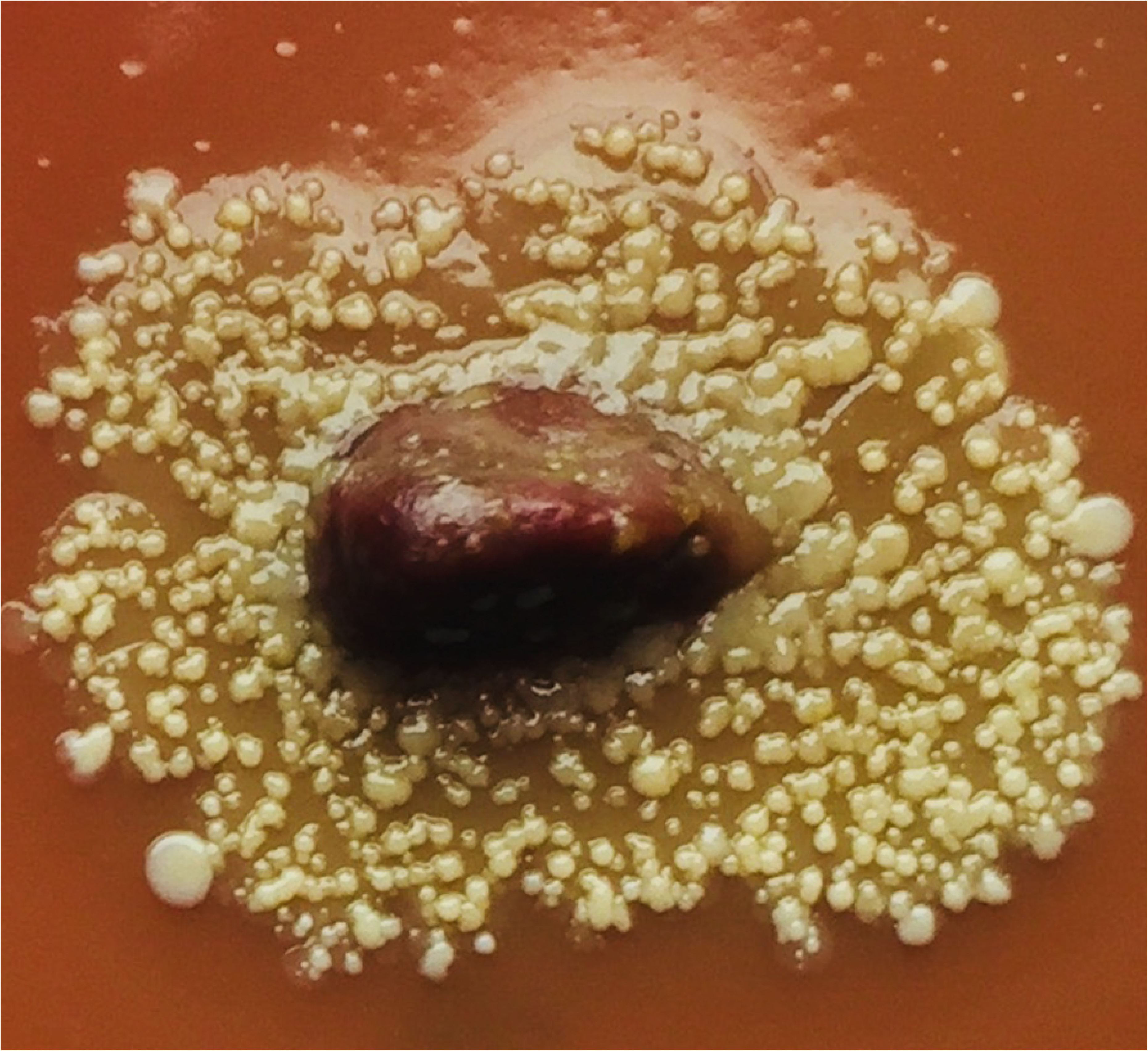
The same specimen as on Fig. 4 one month after on-plate culturing. Multiple separated colonies, some of them increasing in size and spreading from the periphery.

**Fig 5.**
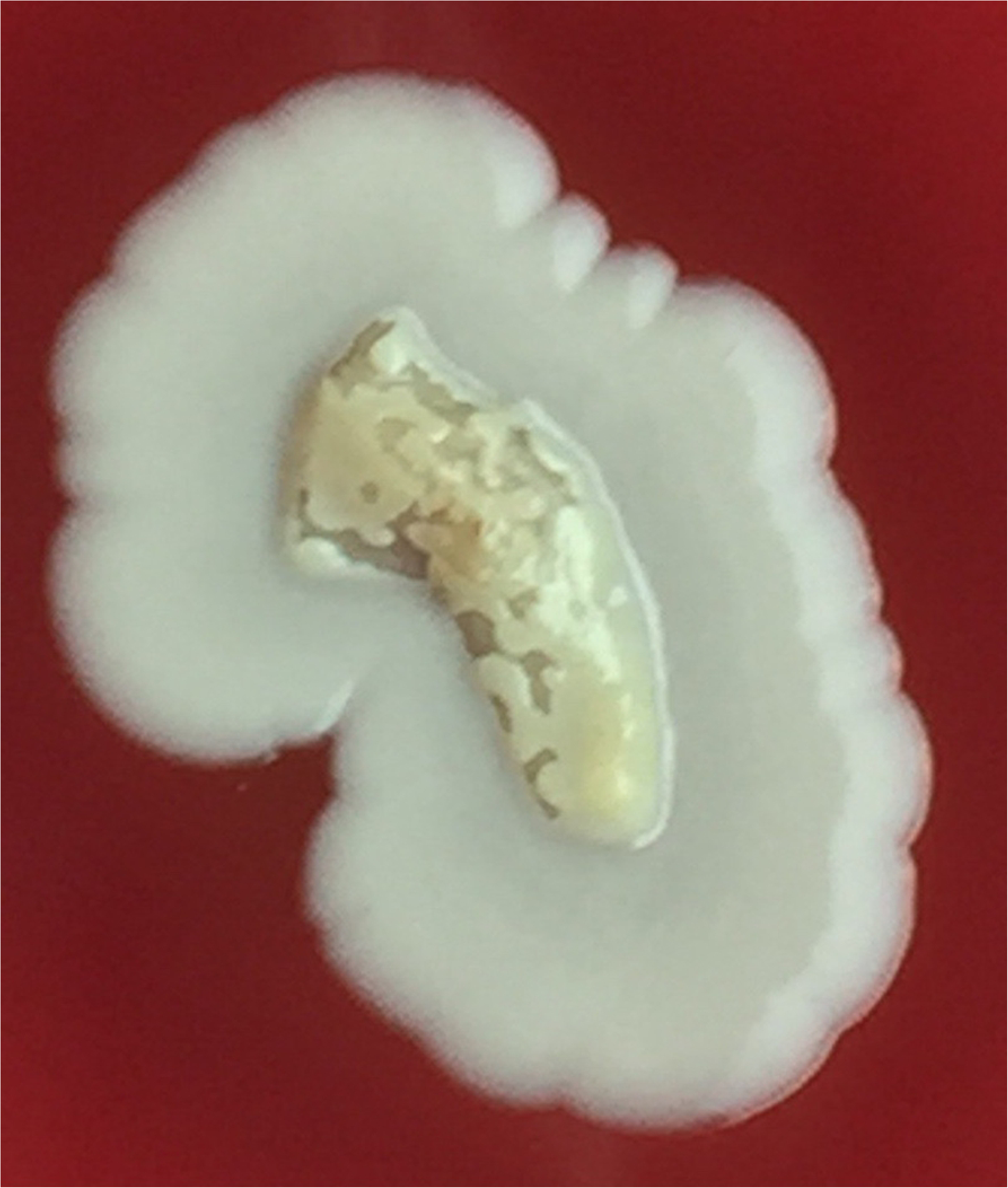
Bacterial colonies seen also on the specimen surface. Picture may indicate that the natural tissue provides nutrients for bacterial proliferation and the culture medium is only secondary to it.

**Fig 6.**
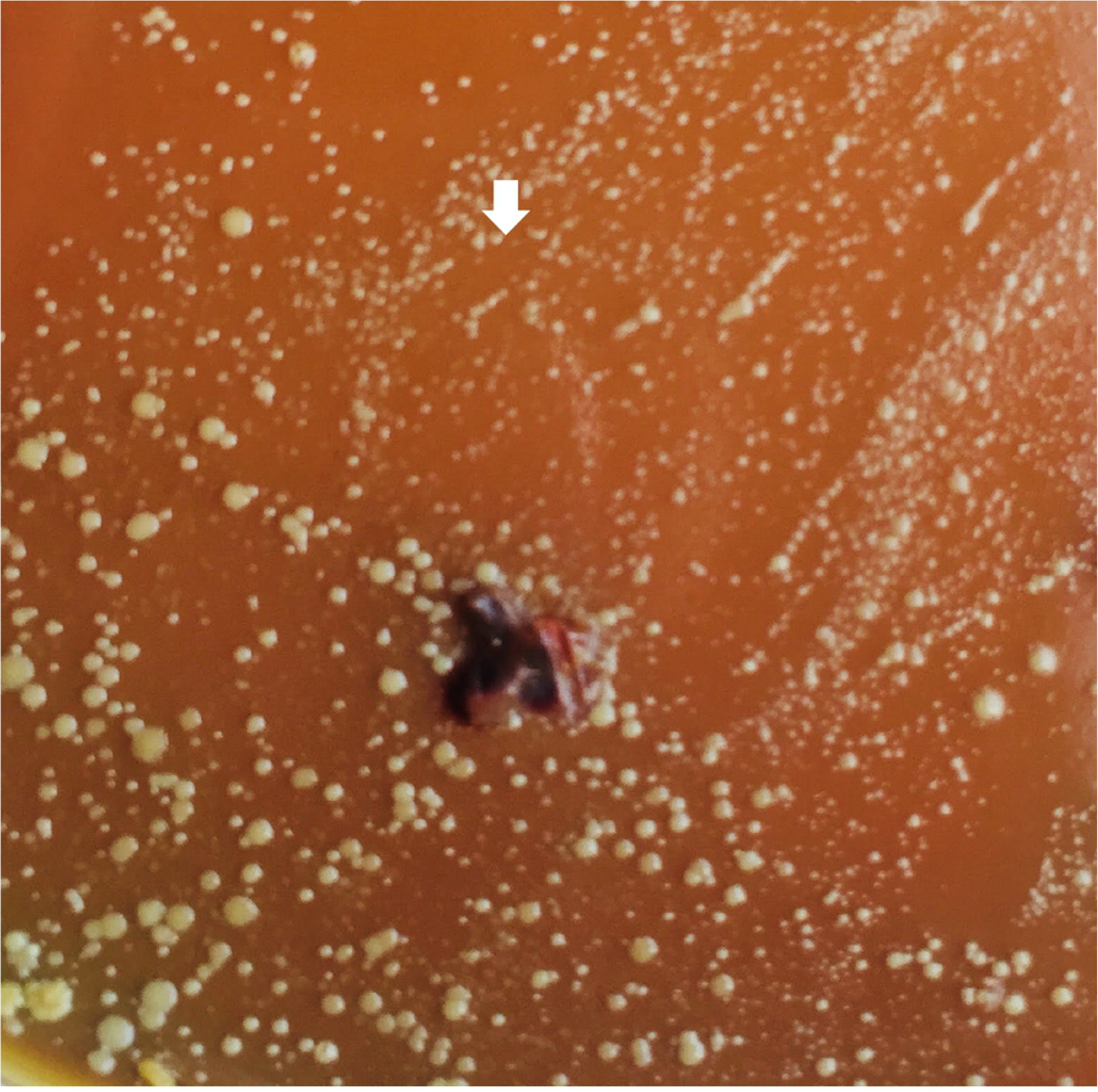
Thyroid specimen cultured on-plate for 4 weeks. Liquid medium was spread upon the plate on week 3 and caused spread of colonies. Note the small variant colonies (arrow). These were colonies of Staphylococcus epidermidis presumably of the “persister” type.

The main identified bacterial phenotype was of coagulase-negative Staphylococci, among them Staphylococcus epidermidis was present in 60% of isolates (Table 1). Less frequently were cultured other coagulase-negative phenotypes. Occasionally, detected were Micrococcus luteus, Aerococcus viridans and Acinetobacter calcoaceticus var lwoffii. The small and also the non-proliferating long term colonies were of Staphylococcus epidermidis.

### Oral vestibule gingival swabs

In 15 consecutive patients directly before thyroid surgery the bacteriological identification from gingiva revealed presence of Staphylococcus aureus and epidermidis in 40% isolates.

### Sensitivity to antibiotics

Antibiotic sensitivity of thyroid and oral isolated coagulase-negative Staphylococci (%) is shown in Table 2. Practically all bacteria responded to the basic antibiotics with the lowest response to penicillin and tetracycline.

**Table 2.** Antibiotic sensitivity patterns of thyroid and oral vestibule Staphylococci coagulase-negative species isolated from thyroid tissue samples and oral cavity swabs (n=10)

|  | Thyroid tissue samples (%) | Oral vestibule swabs (%) |
| --- | --- | --- |
| Penicillin | 60 | 60 |
| Oxacillin coagulase (-) | 100 | 100 |
| Oxacillin coagulase (+) | 100 | 100 |
| Kanamycin | 80 | 80 |
| Tobramycin | 80 | 100 |
| Gentamicin | 100 | 100 |
| Erythromycin | 60 | 60 |
| Lincomycin | 80 | 100 |
| Clindamycin | 80 | 100 |
| Pristinamycin | 100 | 100 |
| Quinupristin | 100 | 100 |
| Tetracycline | 60 | 60 |
| Minocycline | 100 | 100 |
| Levofloxacin | 100 | 100 |
| Ofloxacin | 100 | 100 |
| Linezolid | 100 | 100 |
| Fusidic acid | 80 | 100 |
| Rifampicin | 100 | 100 |
| Fosfomycin | 100 | 80 |
| Nitrofurantoin | 100 | 100 |
| Co-Trimoxazole | 100 | 100 |
| Vancomycin | 100 | 100 |
| Teicoplanin | 100 | 100 |

### Polymerase Chain Reaction Melting Profiles

The DNA electrophoretic patterns of Staphylococcus epidermidis isolated from all sources in oral cavity and thyroid were compared. The similarity between oropharyngeal / nasal cavity and thyroid was from 55 to 88%. Data are shown in Table 3. Representative electrophoresis gel picture is shown on Figure 7.

**Tab 3.** Similarity matrix values on RAPD data among bacterial isolates from ^a)^ oropharyngeal / nasal cavity and ^b)^ thyroid

|  | <sup>a)</sup> Gingival pocket | <sup>a)</sup> Throat | <sup>a)</sup> Oral vestibule | <sup>a)</sup> Nasal cavity | <sup>b)</sup> Thyroid vein | <sup>b)</sup> Thyroid artery | <sup>b)</sup> Thyroid tissue | <sup>b)</sup> Perithyroidal lymph node |
| --- | --- | --- | --- | --- | --- | --- | --- | --- |
| <sup>a)</sup> Gingival pocket | 1,000 |  |  |  |  |  |  |  |
| <sup>a)</sup> Throat | 0,775 | 1,000 |  |  |  |  |  |  |
| <sup>a)</sup> Oral vestibule | 0,723 | 0,828 | 1,000 |  |  |  |  |  |
| <sup>a)</sup> Nasal cavity | 0,729 | 0,939 | 0,889 | 1,000 |  |  |  |  |
| <sup>b)</sup> Thyroid vein | **0,882** | **0,788** | **0,790** | **0,817** | 1,000 |  |  |  |
| <sup>b)</sup> Thyroid artery | **0,870** | **0,788** | **0,783** | **0,819** | 0,963 | 1,000 |  |  |
| <sup>b)</sup> Thyroid tissue | **0,828** | **0,707** | **0,546** | **0,721** | 0,867 | 0,862 | 1,000 |  |
| <sup>b)</sup> Perithyroidal lymph node | **0,714** | **0,644** | **0,745** | **0,768** | 0,945 | 0,940 | 0,764 | 1,000 |

Bold text – similarity of bacterial isolates from oropharyngeal / nasal cavity vs thyroid

**Fig 7.**
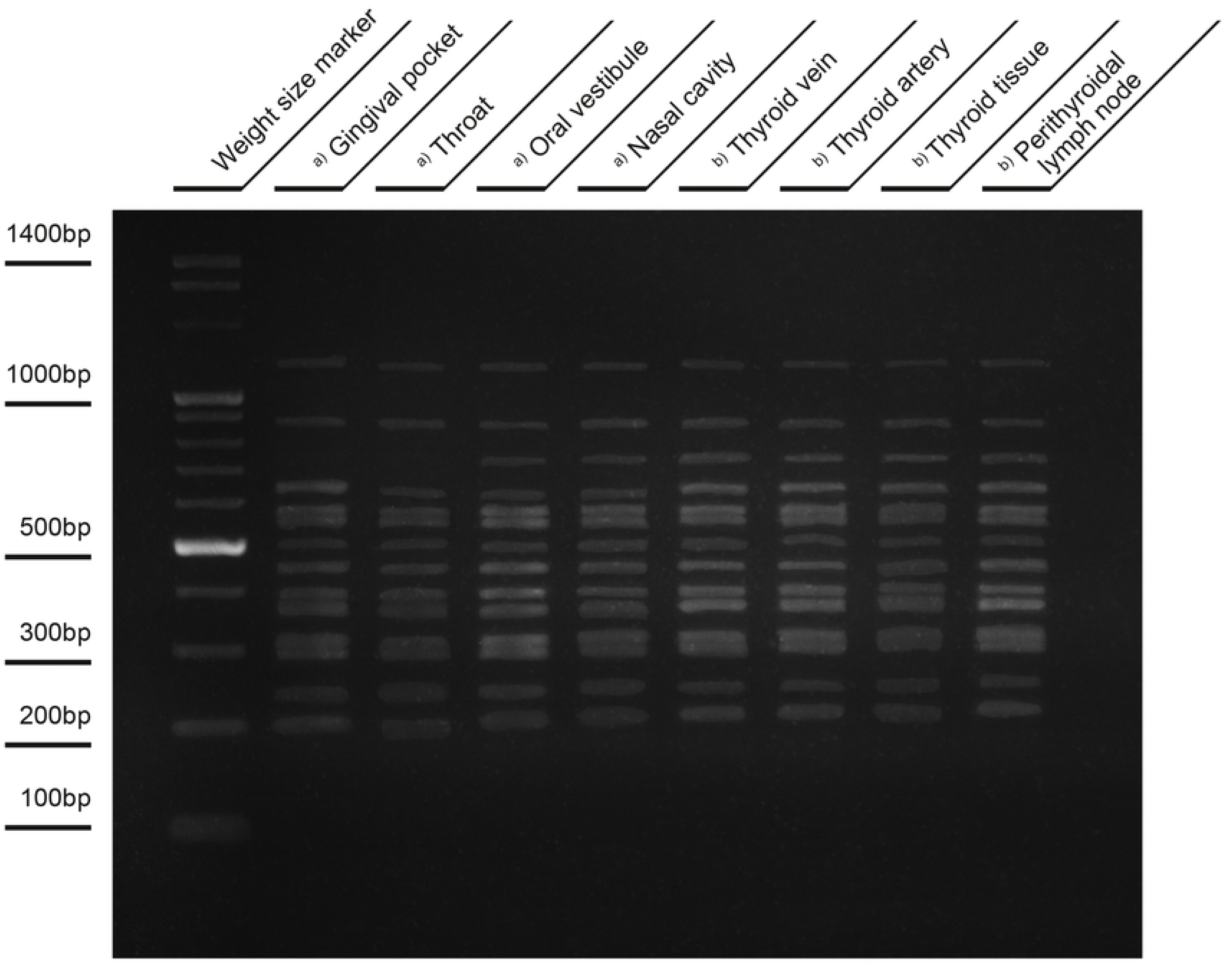
PCR MP of bacterial isolates. a) oropharyngeal / nasal cavity and b) thyroid.

## Discussion

This study provided the following information: a) presence of coagulase-negative Staphylococci in over 50% of goiter parenchyma, vein, artery and adjacent lymphoid tissue specimens shown on the on-plate culture, b) tissue originating colony-forming bacteria appeared on plates on day 3 of culture, in some even as late as after 12-21 days, c) all isolates were sensitive to the basic antibiotics, d) bacterial thyroid and oral DNA tests showed similarities indicating possibility of the oral origin, e) the on-plate time-prolonged cultures showed shrinking of the colonies and upon adding liquid medium formed the small variant colonies. The fact that bacteria did not get into the tested tissue samples from the theatre air or instruments was lack of control bacterial growth. Taken together the main, so far not expected finding, was the overall presence of Staphylococci in the gland not revealing any acute inflammation.

Detection and identification of bacteria colonizing deep tissues remains a difficult task. Microorganisms hidden in tissue niches may not be culturable using the conventional methods. In such cases special methods for their detection become indispensable [23]. We placed the collected tissue material on Columbia agar with sheep blood plate and cultured it for up to 30 days. In this method bacteria present in the tissue slowly proliferate in their in vivo transferred to ex-vivo environment, crawl out and form the on-plate colonies. It enables detection of single bacteria usually difficult to be done in a standard planktonic culture. Bacteria forming colonies moved from the tissue and were transferred to transport media for further close identification. It is our routinely applied method described previously [15]. Using this method we could observe the “dormant” bacteria, forming small colony variants and also large long term culture non-proliferating colonies.

The colonization of thyroid tissues by bacteria could take place long before they were identified. To enable living in a settled unfavorable place for such a long time, bacteria must avoid destruction and develop resistance against host defense, iodine and hydrogen peroxide etc. It is well known that some viable bacteria can be present in tissue but they are non culturable in conventional methods (VBNC) [24]. Microbial dormancy is a widespread phenomenon employed by bacteria to evade environmental threats. Such bacteria retain their basic metabolic activity with slowed vital functions. Following a favorable change in environmental conditions, like in laboratory incubator, placed on Columbia agar plate, dormant forms of bacteria can return to the culturable forms by so-called resuscitation phenomenon, then a small number of surviving cells allows to reproduce the initial population [25–26]. This was not studied in the present investigation.

Detection of microbial cells in thyroid deep tissue of patients without any symptoms of infection may prove the primary or secondary role of bacteria in thyroiditis or other diseases like autoimmune or even cancer formation [27]. Are microbes transported from the primary source to the thyroid via blood as from the gut or veins and lymphatics draining mouth and trachea? The lymph vessels from the gingiva and teeth drain into the submandibular and submental lymph nodes or directly into the deep cervical nodes. So, thyroid gland is drained by lymphatics merging with those draining mouth and nasopharynx tissues. In case of dental or oropharyngeal bacterial infection and subsequent inflammation of the lymphatic system, bacteria present in the lymph may easily reach thyroid gland. Also the venous drainage to the internal jugular vein both from mouth and thyroid gland may facilitate bacterial transport from mouth and subsequent retention of bacteria in the thyroid gland. Patients with Hashimoto thyroid had a higher number of total cervical lymph nodes than the control group, most notably in cervical levels III and IV [28].

The presence of bacteria we detected in the thyroid does not necessarily mean that following the injury they will switch from dormant state to an active state and trigger a massive host response. The presence of bacteria in our body is not only limited to the gastrointestinal tract, respiratory system, urogenital system and exudative glandular organs like the breast gland. The presence of cryptic microbes has been also widely documented in animal healthy deep tissues [29–31]. They can also play a significant role in the pathological processes. Cryptic bacteria of lower limb deep tissues may be cause of inflammatory and necrotic changes in ischemia, venous stasis with varicose veins, and lymphedema [15]. Dormant bacteria were identified in callus of closed fractures of the femur and tibia [16]. Specimens of thrombotic fragments of saphenous vein obtained from limbs without ulcer revealed presence of bacteria [32]. Harvesting of normal great saphenous vein for aortocoronary bypass graft is frequently complicated by infection and delayed wound healing, although taken under strictly aseptic conditions [33–35]. The presence of some oral microbiome bacterial species were shown in clinically non-atherosclerotic coronary and femoral arteries [13].

In the described study the microorganisms found in the thyroid could contribute to the primary thyroid tissue changes. The subject may be controversial as some reports indicate the presence of coagulase-negative Staphylococci can have a beneficial effect on host. Specific strains of Staphylococcus epidermidis can influence host function and have been shown to produce proteins that work together with endogenous host antimicrobial peptides (AMPs) to provide direct protection against Staphylococcus aureus [36,37]. This species are able to influence skin immune function by diminishing inflammation after injury [38].Strains of Staphylococcus epidermidis produce 6-N-hydroxyaminopurine (6-HAP), a molecule that inhibits DNA polymerase activity. 6-HAP selectively inhibits proliferation of tumor lines in vitro and suppresses the growth of melanoma and in vivo [39].

Taken together, our studies showed that thyroid gland deep tissues contain bacteria belonging probably to the persister or most likely VBNC (viable bacteria non-culturable) cells. More than 88% similarity of the genetic pattern of Staphylococcus epidermidis strain from tooth, oropharyngeal and thyroid tissues, estimated with PCR MP technique, suggested their periodontium origin. We did not provide proves for their pathogenicity, nevertheless, their presence in such a high number of specimens should be seriously considered as a possible factor in the pathogenesis of chronic thyroiditis and thyroidectomy wound healing problems. Limitations of the study was lack of evidence of the pathogenicity of the thyroid tissue isolates taking into account their high numerical frequency and no laboratory testing of the revival capacity of the “dormant” Staphylococci found in the gland tissue as well as their in vitro effect on thyroid cell hormone synthesis.

